# Subcortical brain segmentation in 5-year-old children: validation of FSL-FIRST and FreeSurfer against manual segmentation

**DOI:** 10.1101/2021.05.28.445926

**Authors:** Kristian Lidauer, Elmo P. Pulli, Anni Copeland, Eero Silver, Venla Kumpulainen, Niloofar Hashempour, Harri Merisaari, Jani Saunavaara, Riitta Parkkola, Tuire Lähdesmäki, Ekaterina Saukko, Saara Nolvi, Eeva-Leena Kataja, Linnea Karlsson, Hasse Karlsson, Jetro J. Tuulari

**Affiliations:** The FinnBrain Birth Cohort Study, Turku Brain and Mind Center, Department of Clinical Medicine, University of Turku, Finland; Department of Psychiatry, Turku University Hospital, University of Turku, Turku, Finland; Turku Collegium for Science, Medicine and Technology, University of Turku, Turku, Finland; Department of Psychiatry, University of Oxford, UK (Sigrid Juselius Fellowship), United Kingdom; Department of Radiology, University of Turku, Turku, Finland; Department of Radiology, Turku University Hospital, Turku, Finland; Turku Institute for Advanced Studies, University of Turku, Turku, Finland; Department of Psychology and Speech-Language Pathology, University of Turku, Turku, Finland; Department of Medical Physics, Turku University Hospital, Turku, Finland; Centre for Population Health Research, University of Turku and Turku University Hospital, Turku, Finland; Department of Paediatric Neurology, Turku University Hospital and University of Turku, Turku, Finland

**Keywords:** brain, child, neuroimaging, brain (growth & development)

## Abstract

Developing accurate subcortical volumetric quantification tools is crucial for neurodevelopmental studies, as they could reduce the need for challenging and time-consuming manual segmentation. In this study the accuracy of two automated segmentation tools, FSL-FIRST (with three different boundary correction settings) and FreeSurfer were compared against manual segmentation of subcortical nuclei, including the hippocampus, amygdala, thalamus, putamen, globus pallidus, caudate and nucleus accumbens, using volumetric and correlation analyses in 80 5-year-olds.

Both FSL-FIRST and FreeSurfer overestimated the volume on all structures except the caudate, and the accuracy varied depending on the structure. Small structures such as the amygdala and nucleus accumbens, which are visually difficult to distinguish, produced significant overestimations and weaker correlations with all automated methods. Larger and more readily distinguishable structures such as the caudate and putamen produced notably lower overestimations and stronger correlations. Overall, the segmentations performed by FSL-FIRST’s Default pipeline were the most accurate, while FreeSurfer’s results were weaker across the structures.

In line with prior studies, the accuracy of automated segmentation tools was imperfect with respect to manually defined structures. However, apart from amygdala and nucleus accumbens, FSL-FIRST’s agreement could be considered satisfactory (Pearson correlation > 0.74, Intraclass correlation coefficient (ICC) > 0.68 and Dice Score coefficient (DSC) > 0.87) with highest values for the striatal structures (putamen, globus pallidus and caudate) (Pearson correlation > 0.77, ICC > 0.87 and DSC > 0.88, respectively). Overall, automated segmentation tools do not always provide satisfactory results, and careful visual inspection of the automated segmentations is strongly advised.

## Introduction

The subcortical structures of the brain are responsible for numerous important functions and their development can be affected by early life environmental exposures and experiences (Lee et al., 2019; Pulli et al., 2019). Therefore, it is crucial to gather accurate information about them in magnetic resonance imaging (MRI) studies conducted in paediatric populations. Accurate segmentation of paediatric MR images is challenging, partly due to the variation in pre-processing and segmentation protocols (Hashempour et al., 2019; Schoemaker et al., 2016). Several segmentation protocols have been developed for adult brains, but they cannot be directly applied in segmenting children brain since children MR images have different contrast and comparatively lower resolution than adults’ images (Gousias et al., 2012; Moore et al., 2014; Morey et al., 2009). Manual segmentation is currently considered the gold standard in volumetric segmentation. While it is considered the most accurate method, it is highly time consuming and requires expertise for adequate results. Furthermore, a major downside is the subjective approach in estimating the shapes and sizes of the structures, which may cause reproducibility issues that may be even more pronounced in larger samples.

Several software have been developed for automated segmentation of the brain. In this study, we focused on two mainstream analysis pipelines. One is FSL-FIRST from the FMRIB software library (https://fsl.fmrib.ox.ac.uk/fsl/fslwiki/FIRST). FSL-FIRST is a segmentation tool that uses the template based on manually segmented images to construct the shape of the automated segmentation models. It utilises the Active Appearance Model (AAM) combined with a Bayesian framework, which allows probabilistic relationships between voxel intensity and the shapes of different structures (Patenaude et al., 2011). The other is FreeSurfer (https://surfer.nmr.mgh.harvard.edu/), which is an open-source software suite for processing and analysing MR images. FreeSurfer uses a five-stage volume-based stream for segmenting subcortical structures. Final segmentation is based on a subject-independent probabilistic atlas and subject specific values. Both FSL-FIRST and FreeSurfer use a training dataset for the basis of segmentation and utilise probabilistic computing to determine the final shape and volume of each structure. Although both FSL-FIRST and FreeSurfer were originally developed for adult brain imaging and utilize adult templates, they have also been used in paediatric imaging (Barch et al., 2019; Schoemaker et al., 2016).

Consistent overestimation of subcortical volumes regarding both FreeSurfer and FSL-FIRST (Cherbuin et al., 2009; Doring et al., 2011) has been a common finding in previous studies. This result has been documented in paediatric populations on the hippocampus and amygdala (Mulder et al., 2014; Schoemaker et al., 2016). The study by Schoemaker et al. also found that the consistency between manual segmentation and FreeSurfer was better than between manual segmentation and FSL-FIRST in children aged between 6 and 11 years (Schoemaker et al., 2016). While the reliability of these segmentation methods has been assessed in multiple studies in the medial temporal lobe structures, there has been little research including the striatal structures.

The aim of this study was to compare the accuracy of FSL-FIRST and FreeSurfer against the gold standard manually corrected segmentation on subcortical structures, including the hippocampus, amygdala, thalamus, putamen, globus pallidus (GP), caudate and nucleus accumbens, in paediatric populations. Therefore, we compared the volumes of all the structures extracted from each segmentation method. Furthermore, we analysed the shape of the segmentation models to determine the areas where the automated segmentation tools overestimated or underestimated the size of the structures and their borders. This was a feasibility study that critically assessed the extent to which adult delineation software can be used to segment child brain images that have nearly adult-like contrast pattern in T1-weighted images and are close in size to adult brain.

## Material and Methods

This study was conducted in accordance with the Declaration of Helsinki, and it was approved by the Joint Ethics Committee of the University of Turku and the Hospital District of Southwest Finland (07.08.2018) §330, ETMK: 31/180/2011.

### Subjects

MRI scans were acquired in children as part of the FinnBrain Birth Cohort Study (www.finnbrain.fi), which was started in 2011. The main goal of the cohort is to study the effects of genes and environment on the development and mental health of children (Karlsson et al., 2018). Initial recruitment of FinnBrain Birth Cohort Study was performed systematically in routine ultrasound examinations during the 12^th^ week of gestation. For the 5-year neuroimaging visit, we primarily recruited participants that had a prior visit to neuropsychological measurements at ca. 5 years of age (n = 76). However, there were a few exceptions: three participants were included without a neuropsychological visit, as they had an exposure to maternal prenatal synthetic glucocorticoid treatment (recruited separately for a nested case-control sub study). The data additionally included one participant that was enrolled for a pilot scan. The exclusion criteria for this study were: 1) born before gestational week 35 (born before gestational week 32 in the synthetic glucocorticoid treatment group), 2) developmental anomaly or abnormalities in senses or communication (e.g. congenital heart disease, blindness, deafness), 3) known long-term medical diagnosis (e.g. epilepsy, autism, attention deficit hyperactivity disorder (ADHD)), 4) ongoing medical examinations or clinical follow up in a hospital (meaning there has been a referral from primary care setting to experts), 5) child use of continuous, daily medication (including per oral medications, topical creams and inhalants. One exception to this was desmopressin (®Mirinin) medication, which was allowed), 6) history of head trauma (defined as concussion necessitating clinical follow up in a health care setting or worse), 7) metallic ear tubes (to assure good-quality scans), and routine MRI contraindications.

In this study we used a representative subsample of 80 T1-weighted brain images, which were all visually inspected by a single expert rater (Kristian Lidauer). The sample included 34 girls and 46 boys aged between 5 and 5.5 years (mean age 5.34 years, SD = 0.06).

### Study visit

The subjects were recruited for the neuroimaging visits via phone calls by a research staff member. On the first call the families were given general information about the study and the inclusion and exclusion criteria were checked. The follow-up call was made to confirm the participation, and we gave instructions to practice for the MRI visit at home. A member of the research staff made a home visit before the scan to deliver earplugs and headphones, to give more detailed information about the visit, and to answer any remaining questions. An added benefit of the home visit was the chance to meet the participating child and that way start the familiarization with the research staff, which helped the preparations on the scanning day. A written consent was acquired from both parents before the MRI scan as well as verbal assent from the child.

Multiple methods were applied to reduce anxiety and make the visit feel as safe as possible (many of the methods have been described in earlier studies (Greene, Black, & Schlaggar, 2016). The visit was conducted in a child-friendly manner with a flexible timetable in the preparation before the scan, and we did our best to accommodate in order to befit the child in cooperation with the family. The participants were scanned awake. During the structural imaging the subjects were allowed to watch a cartoon or a movie of their choice. A parent and a research staff member were present in the scanner room throughout the scan. Everyone in the room had their hearing protected with earplugs and headphones. The maximum scan time was 60 minutes, and the subjects were allowed to stop the scan at any time. For a more detailed description of the study visits, please see (Pulli et al., 2021) and (Copeland et al., 2021).

### MRI acquisition

Participants were scanned using a Siemens Magnetom Skyra fit 3T with a 20-element head/neck matrix coil. We used Generalized Autocalibrating Partially Parallel Acquisition (GRAPPA) technique to accelerate image acquisition (parallel acquisition technique [PAT] factor of 2 was used). For the purposes of the current study, we acquired a high resolution three-dimensional T1-weighted magnetization prepared rapid acquisition gradient-echo sequence (MPRAGE) in sagittal plane with the following sequence parameters: TR = 1900 ms, TE = 3.26 ms, TI = 900 ms, FA = 9 degrees, voxel size = 1.0×1.0×1.0 mm³, FOV = 256 mm. In addition, the max. 60-minute scanning protocol included a T2 turbo spin echo (TSE), a 7-minute resting state functional MRI, and a DTI-sequence. The T1 scans were planned as per recommendations of the FreeSurfer developers (https://surfer.nmr.mgh.harvard.edu/fswiki/FreeSurferWiki?action=AttachFile&do=get&target=FreeSurfer_Suggested_Morphometry_Protocols.pdf, at the time of writing).

### Automated segmentation of the subcortical nuclei using FSL-FIRST

The automated segmentation of the subcortical structures was performed using FSL-FIRST 5.0.9 (https://fsl.fmrib.ox.ac.uk/fsl/fslwiki/FIRST), a freely available automated segmentation tool provided by the FMRIB software library. FSL-FIRST uses a training data-based approach combined with a Bayesian probabilistic model to determine the most probable shape of the structure given the intensities of the T1-image. More detailed information about the technical process can found in an article by Patenaude et al. (Patenaude et al., 2011). In this study, we segmented the T1-images using FSL-FIRST with three different boundary correction settings. The *FSL Default* method uses different options based on empirical observations for each different structure. The *FSL Fast* option uses an FSL-FAST based tissue-type classification to determine the final shape of the model. For the third boundary correction option we chose *FSL None*, which does not use any boundary correction settings. After running the pipelines, a voxel count was performed to estimate the volumes produced by each different method.

### Automated segmentation of the subcortical nuclei using FreeSurfer

The other automated segmentation software used in this study was FreeSurfer 6.0 (https://surfer.nmr.mgh.harvard.edu/), a freely available open software neuroimage analysis suite. We used the recon-all pipeline with default settings consisting of several stages. In brief, the process includes motion correcting and averaging of multiple T1 images, which is proceeded by removal of non-brain tissue using a watershed/surface deformation procedure, after which the images are transferred into a Talairach space, where the white matter and subcortical grey matter are segmented by labelling each voxel based on the probabilities from a manually edited training dataset and the intensities of the T1-image. The technical details of the FreeSurfer process are described more in-depth in prior publications (Fischl et al., 2002, 2004; Segonne et al., 2004). The volumes were extracted with “asegstats2table” command.

### Manual segmentation of the subcortical nuclei

Manual segmentation was done by editing the models produced by *FSL None*. We visually inspected the results of all three FSL-FIRST pipelines and chose *FSL None*, because it required the least amount of editing. The subcortical structures were segmented by a single expert rater (Kristian Lidauer) using the software FslView (https://fsl.fmrib.ox.ac.uk/fsl/fslwiki/FslView). The rater was experienced in manual segmentation of infant brain MR images and templates (Acosta et al., 2020; Hashempour et al., 2019) across a period of two years before starting the current study (2018-2020).

The use of initial estimates from FSL-FIRST significantly reduced the working time as compared to full manual segmentation. It also made the work easier as the main task for the investigators was correction of the borders. This process was guided by prior work for striatal structures (Perlaki et al., 2017) and the thalamus (Owens-Walton et al., 2019; Power et al., 2015) as well as our prior work for amygdala and hippocampus segmentation, which is provided in our recent open access article (Hashempour et al., 2019).

The manual edits were performed on “initial estimates” that saved time. The edits were documented on 40 randomly chosen subjects of the total 80 to highlight important areas for quality control. The anatomical delineations that we incorporated into locally adapted procedures are in line with prior work (de Macedo Rodrigues et al., 2015). Manual segmentations/edits were performed in a slice-by-slice manner to carefully trace the correct anatomical border and reviewed in axial, coronal and sagittal planes for a three-dimensional consistency of the segmentations. Finally, all segmentations were checked for accuracy by senior scientist (Jetro J. Tuulari). The accuracy check was performed with fsleyes and entailed: 1) selection of a reference segmentation with all structures accurately delineated, 2) opening three segmentations at a time and comparing them against the reference segmentation, 3) checking bilateral structures from each one by browsing the structure in all 3D planes and checking the borders with intermittent opening and closing the overlay to check the consistency of the borders. This process took about 15 minutes per three segmentation (ca. 7 hours in the final round of quality control).

A voxel count was then concluded with fslmaths to estimate the volumes of the manually segmented structures.

### Statistical analysis

All statistical analyses and plotting of the results were performed using R tools v.4.0 (https://www.r-project.org/) and R-Studio 1.3 (https://rstudio.com/). For the plots and following analyses we used irr, ggplot2, gridExtra, grid and gtable libraries.

The volumetric difference between automated segmentation and manual segmentation was calculated as the percentage using the following equation (Schoemaker et al., 2016): %VD = [(V_a_ – V_m_)/V_m_] * 100%, where V_a_ is the automated volume and V_m_ is the manually segmented volume. A negative result indicates that the automated method underestimated the volume whereas a positive value shows that the automated method overestimated the volume.

Pearson correlations were calculated to measure the strength of the association between manual segmentation and the different automated techniques for each individual structure. A strong correlation would indicate good consistency between methods. To estimate reproducibility between different techniques and estimation bias, we computed intraclass correlation coefficients (ICC). We used a two-way mixed effect model with absolute agreement and average measures (ICC type A, k) as specified by McGraw and Wong (McGraw & Wong, 1996), which is a model not defined in the commonly used Shrout and Fleiss convention (Shrout & Fleiss, 1979). A high value would confirm a good reproducibility between two raters. There are no fixed guidelines on how to interpreter ICC values, but in previous studies a coefficient of 0.70 has been considered as the minimum for establishing an adequate reliability between two raters (Terwee et al., 2007).

To determine the spatial overlap of the structures we conducted Dice score coefficient (DSC) analysis between manual and automated segmentation methods. The value of DSC ranges from 0, indicating no spatial overlap between structures, to 1, indicating complete overlap (Zou et al., 2004).

## Results

### Volumetric differences between FSL-FIRST pipelines

*FSL None* produced the highest volumes for the hippocampus, amygdala, caudate and nucleus accumbens, and produced the same result as the *FSL Default* pipeline in the other three structures: the putamen, GP and the thalamus. The mean volumes for the left and right hippocampus were 4244.95 (SD = 575.67) and 4434.70 (SD = 531.64), respectively, and for the left and right amygdala 1377.63 (SD = 232.26) and 1306.54 (SD = 228.94), respectively. The other pipelines, *FSL Default* and *FSL Fast*, had considerably lower volumes for the hippocampus and amygdala and yielded the exact same result for both structures. The mean volumes for the left and right hippocampus were 3412.41 (SD = 441.28) and 3551.45 (SD = 415.35), respectively, and for the left and right amygdala 1096.85 (SD = 203.91) and 1053.94 (SD = 194.49), respectively. *FSL Default* and *FSL Fast* performed very similarly throughout and showed the exact same volumes also for the caudate and the nucleus accumbens. The volumetric unit used is 1 voxel = 1 mm^3.^ The volumes for each pipeline and structure are presented in **Table 1**. The identical results in some of the structures are caused by utilizing the same boundary correction options.

**Table 1.**
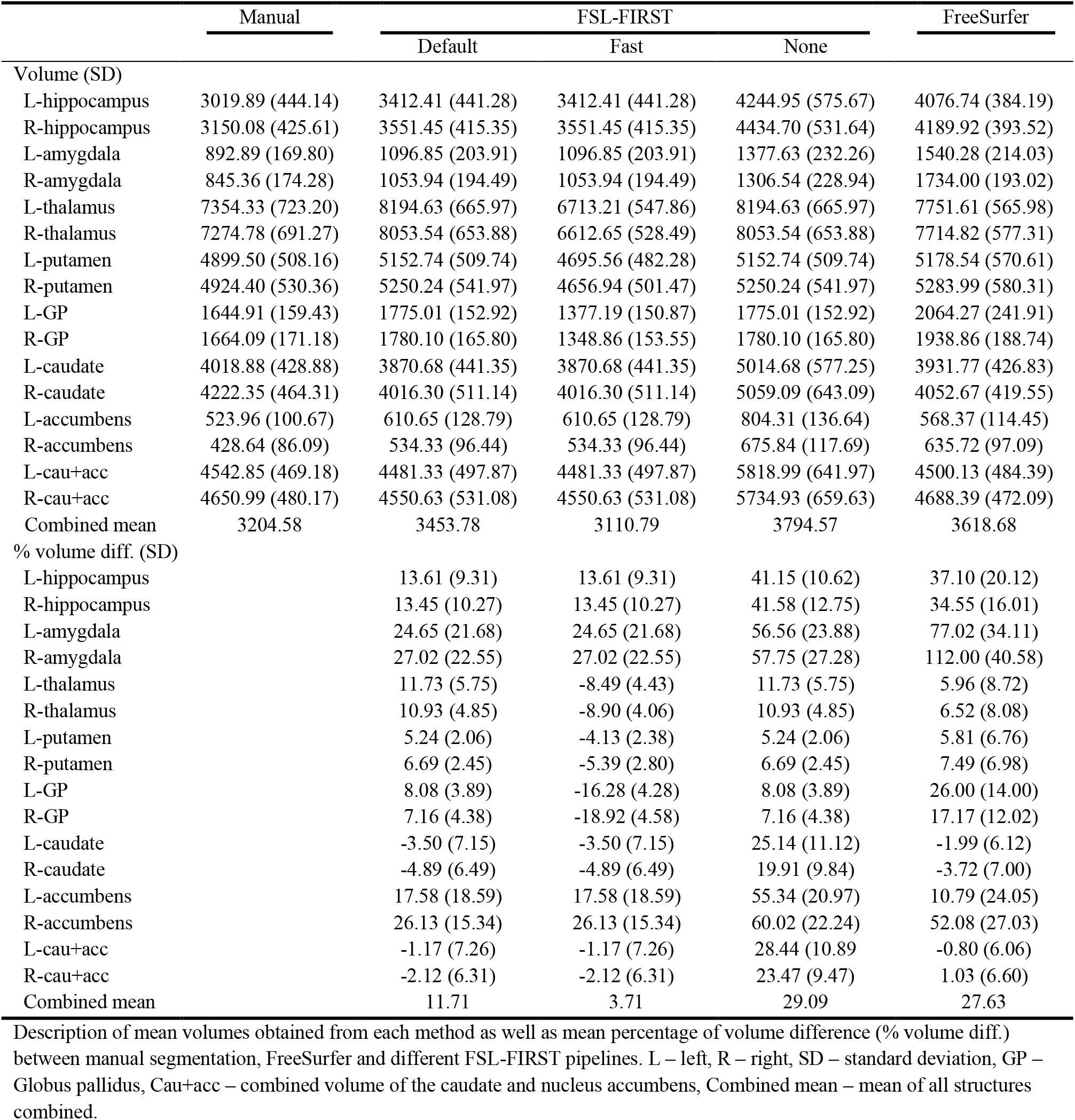
Comparison of mean (standard deviation) volumes and percentage of volume difference between techniques. The volumetric unit used is 1 voxel (= 1 mm³).

The volume difference between FSL-FIRST and manual segmentation was highest with the *FSL None* pipeline. *FSL None* overestimated volumes for the left and right hippocampus 41.15% (SD = 10.62) and 41.58% (SD = 12.75), respectively, and for the left and right amygdala 56.56% (SD = 23.88) and 57.75% (SD = 27.28), respectively. *FSL Default* and *FSL Fast* overestimated volumes less, for the left and right hippocampus 13.61% (SD = 9.31) and 13.45% (SD = 10.27), respectively, and for the left and right amygdala 24.65% (SD = 21.68) and 27.02% (SD = 22.55), respectively, for both pipelines. *FSL Fast* underestimated volumes for the putamen, GP, thalamus and caudate, while *FSL Default* underestimated the caudate volume. *FSL None* overestimated the volume for every structure. The percentage differences for each structure and each pipeline are presented in **Table 1**.

### FSL-FIRST volumetric correlation analysis

Pearson correlation coefficient between *FSL None* and manual segmentation for the left and right hippocampus was r = 0.86 and r = 0.75, respectively, and r = 0.67 for both the left and right amygdala. The correlation for *FSL Default* and *FSL Fast* pipelines for the left and right hippocampus was r = 0.83 and r = 0.74, respectively, and for the left and right amygdala r = 0.61 and r = 0.66, respectively. For the other structures, all three pipelines had similar correlation values, which are all presented in **Table 2**. A scatter plot illustration for all structures and methods is provided in **Figure 1**.

**Table 2.** Comparison of correlation analysis between manual and automated segmentation techniques (FSL-FIRST, FreeSurfer).

|  | FSL-FIRST |  |  | FreeSurfer |
| --- | --- | --- | --- | --- |
|  | Default | Fast | None |  |
| PCC |  |  |  |  |
| L-hippocampus | 0.83 | 0.83 | 0.86 | 0.47 |
| R-hippocampus | 0.74 | 0.74 | 0.75 | 0.54 |
| L-amygdala | 0.61 | 0.61 | 0.67 | 0.34 |
| R-amygdala | 0.66 | 0.66 | 0.67 | 0.47 |
| L-thalamus | 0.86 | 0.87 | 0.86 | 0.60 |
| R-thalamus | 0.89 | 0.88 | 0.89 | 0.61 |
| L-putamen | 0.98 | 0.97 | 0.98 | 0.82 |
| R-putamen | 0.98 | 0.96 | 0.98 | 0.84 |
| L-GP | 0.94 | 0.89 | 0.94 | 0.49 |
| R-GP | 0.92 | 0.87 | 0.92 | 0.52 |
| L-caudate | 0.78 | 0.78 | 0.69 | 0.84 |
| R-caudate | 0.87 | 0.87 | 0.80 | 0.80 |
| L-accumbens | 0.69 | 0.69 | 0.77 | 0.44 |
| R-accumbens | 0.81 | 0.81 | 0.76 | 0.56 |
| L-cau+acc | 0.77 | 0.77 | 0.70 | 0.83 |
| R-cau+acc | 0.85 | 0.85 | 0.78 | 0.81 |
| Combined mean | 0.83 | 0.82 | 0.82 | 0.60 |
| ICC (A,k) |  |  |  |  |
| L-hippocampus | 0.75 | 0.75 | 0.34 | 0.20 |
| R-hippocampus | 0.68 | 0.68 | 0.28 | 0.23 |
| L-amygdala | 0.55 | 0.55 | 0.29 | 0.09 |
| R-amygdala | 0.58 | 0.58 | 0.31 | 0.07 |
| L-thalamus | 0.66 | 0.72 | 0.66 | 0.66 |
| R-thalamus | 0.69 | 0.70 | 0.69 | 0.66 |
| L-putamen | 0.93 | 0.95 | 0.93 | 0.84 |
| R-putamen | 0.90 | 0.92 | 0.90 | 0.82 |
| L-GP | 0.82 | 0.53 | 0.82 | 0.26 |
| R-GP | 0.85 | 0.46 | 0.85 | 0.39 |
| L-caudate | 0.85 | 0.85 | 0.37 | 0.90 |
| R-caudate | 0.89 | 0.89 | 0.53 | 0.85 |
| L-accumbens | 0.69 | 0.69 | 0.33 | 0.58 |
| R-accumbens | 0.65 | 0.65 | 0.31 | 0.27 |
| L-cau+acc | 0.87 | 0.87 | 0.31 | 0.91 |
| R-cau+acc | 0.91 | 0.91 | 0.43 | 0.90 |
| Combined mean | 0.75 | 0.71 | 0.54 | 0.49 |

Pearson correlation coefficients (PCC) and intraclass correlation coefficients (ICC) (A, k) computed between manual and automatic segmentation volumes. L – left, R – right, GP – Globus Pallidus, Cau+acc – combined volume of the caudate and nucleus accumbens, Combined mean – mean of all structures combined.

**Figure 1.**
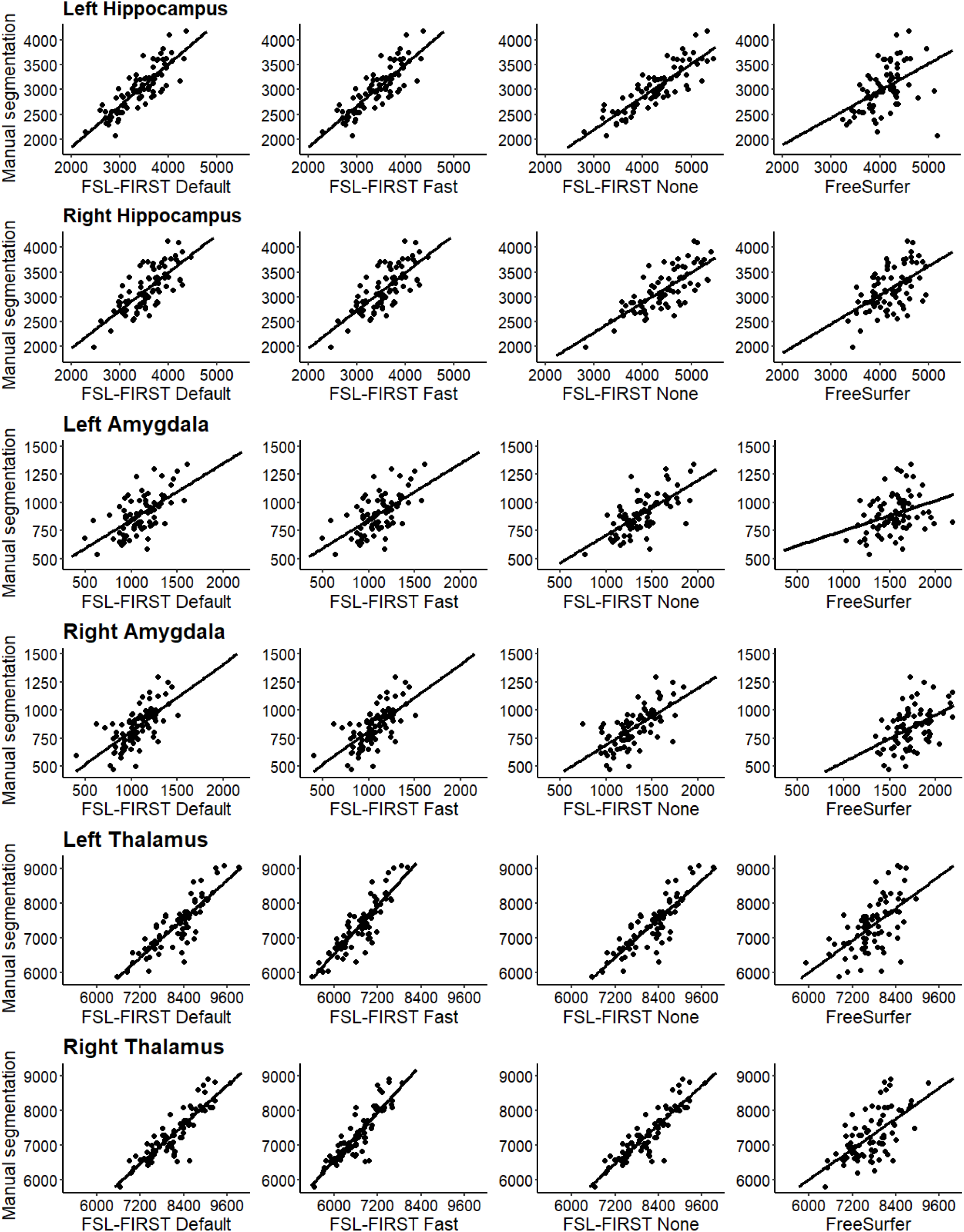
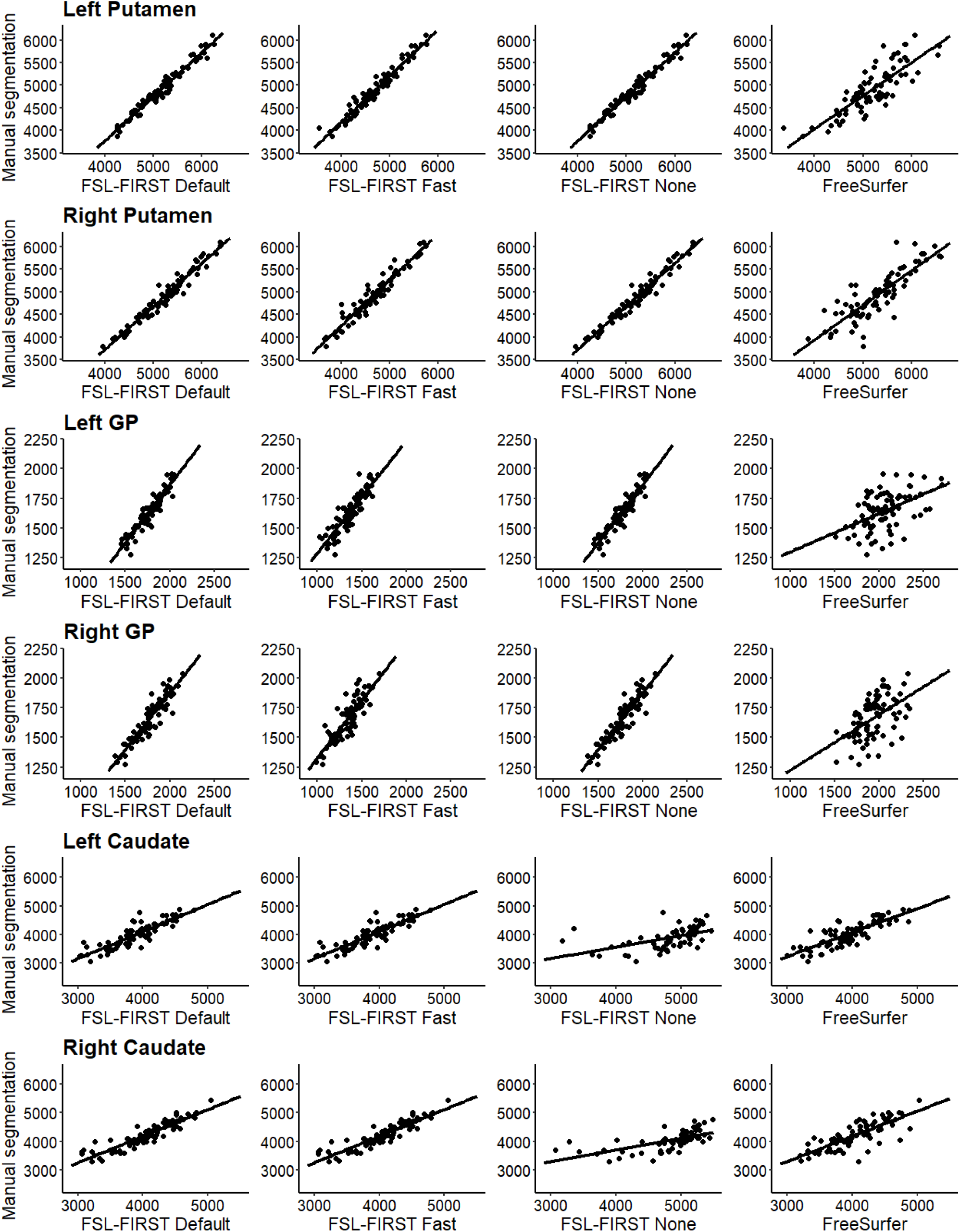
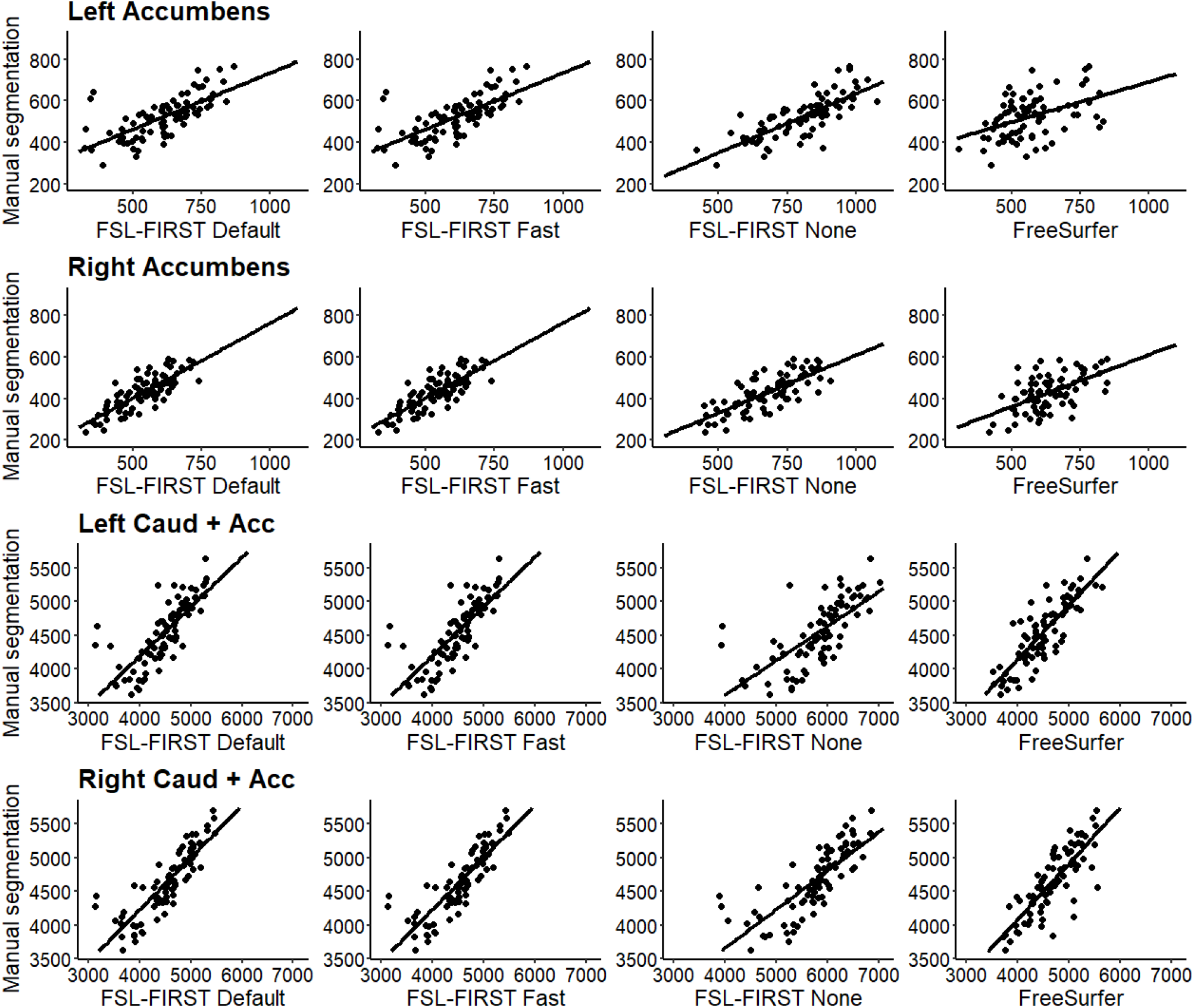
Scatter plots of automated segmentation methods against manual segmentation.

Intraclass correlation coefficient (A, k) between *FSL None* and manual segmentation for the left and right hippocampus was ICC = 0.34 and ICC = 0.28, respectively, and for the left and right amygdala ICC = 0.29 and ICC = 0.31, respectively. The correlations for *FSL Default* and *FSL Fast* pipelines were notably higher than *FSL None’s* for the left and right hippocampus, ICC = 0.75 and ICC = 0.68, respectively and for the left and right amygdala ICC = 0.55 and ICC = 0.58, respectively. Intraclass correlation values for each structure and pipeline are presented in **Table 2**. *FSL None* had the lowest intraclass correlations in almost all structures. *FSL Default* and *FSL Fast* had similar values except for the GP, where *FSL Fast* produced notably lower correlations, ICC = 0.53 and ICC = 0.46 for the left and right GP, respectively, compared to the *FSL Defaults* and *FSL None’s* ICC = 0.82 and ICC = 0.85, for the left and right GP, respectively.

### FreeSurfer volumetric analysis

FreeSurfers volume for the left and right hippocampus was 4076.74 (SD = 384.19) and 4189.92 (SD = 393.52), respectively, and for the left and right amygdala 1540.28 (SD = 214.03) and 1734.00 (SD = 193.02), respectively. FreeSurfer produced the higher volumes than any of the FSL-FIRST pipelines in the amygdala, putamen and GP. Compared to manual segmentation, FreeSurfer had higher volumes in all structures except for the caudate. Percentage difference between FreeSurfer and manual segmentation mean volume for the left and right hippocampus was 37.10% (SD = 20.12) and 34.55% (SD = 16.01), respectively, and for the left and right amygdala 77.02% (SD = 16.01) and 112.00% (SD = 40.58), respectively. Mean volumes and percentage differences for all other structures are presented in **Table 1**.

### FreeSurfer volumetric correlation analysis

Pearson correlation coefficients between FreeSurfer and manual segmentation were lower than any of the FSL-FIRST pipelines in all structures except the caudate, where the values were similar. Pearson correlation coefficient for the left and right hippocampus was r = 0.47 and r = 0.54, respectively, and for the left and right amygdala r = 0.34 and r = 0.47, respectively. Intraclass correlation coefficient (A, k) for the left and right hippocampus was ICC = 0.20 and ICC = 0.23, respectively, and for the left and right amygdala ICC = 0.09 and ICC = 0.07, respectively. FreeSurfer produced overall lower intraclass correlation values except for the caudate, where its values were similar compared to *FSL Default* and *FSL Fast* pipelines, for the left and right caudate ICC = 0.90 and ICC = 0.85, respectively. Pearson and intraclass correlation coefficient values for all structures are presented in **Table 2**.

### Dice score coefficient analysis

Dice score coefficient values between manual segmentation and automated methods were good across the board. FSL-FIRST provided overall slightly higher scores than FreeSurfer. *FSL Default* and *FSL Fast* produced the highest values for the left and right hippocampus, DSC = 0.87 (SD = 0.03) and DSC = 0.88 (SD = 0.03), respectively, for both pipelines. FreeSurfer yielded weaker results for the left and right hippocampus, DSC = 0.76 (SD = 0.05) and DSC = 0.78 (SD = 0.04), respectively. All automated techniques produced lower results for the amygdala than the hippocampus. *FSL Default* and *FSL Fast* had the highest score regarding the left and right amygdala, DSC = 0.73 (SD = 0.05) and DSC = 0.73 (SD = 0.06), respectively. FreeSurfers result for the left and right amygdala was DSC = 0.62 (SD = 0.07) and DSC = 0.60 (SD = 0.07), respectively. Dice score coefficient values for all structures and methods are presented in **Table 3**.

**Table 3.** Comparison of mean Dice score coefficient values between manual and automated segmentation techniques.

|  | FSL-FIRST |  |  | FreeSurfer |
| --- | --- | --- | --- | --- |
|  | Default | Fast | None |  |
| DSC (SD) |  |  |  |  |
| L-hippocampus | 0.87 (0.03) | 0.87 (0.03) | 0.83 (0.04) | 0.76 (0.05) |
| R-hippocampus | 0.88 (0.03) | 0.88 (0.03) | 0.83 (0.04) | 0.78 (0.04) |
| L-amygdala | 0.73 (0.05) | 0.73 (0.05) | 0.72 (0.05) | 0.62 (0.07) |
| R-amygdala | 0.73 (0.06) | 0.73 (0.06) | 0.71 (0.06) | 0.60 (0.07) |
| L-thalamus | 0.95 (0.02) | 0.91 (0.01) | 0.95 (0.02) | 0.88 (0.02) |
| R-thalamus | 0.95 (0.02) | 0.91 (0.01) | 0.95 (0.02) | 0.89 (0.02) |
| L-putamen | 0.98 (0.01) | 0.95 (0.01) | 0.98 (0.01) | 0.86 (0.02) |
| R-putamen | 0.98 (0.01) | 0.95 (0.01) | 0.98 (0.01) | 0.85 (0.03) |
| L-GP | 0.98 (0.01) | 0.88 (0.03) | 0.98 (0.01) | 0.80 (0.05) |
| R-GP | 0.97 (0.02) | 0.87 (0.03) | 0.97 (0.02) | 0.79 (0.06) |
| L-caudate | 0.88 (0.04) | 0.88 (0.04) | 0.86 (0.05) | 0.87 (0.03) |
| R-caudate | 0.90 (0.03) | 0.89 (0.03) | 0.90 (0.04) | 0.87 (0.02) |
| L-accumbens | 0.84 (0.05) | 0.84 (0.05) | 0.84 (0.05) | 0.62 (0.07) |
| R-accumbens | 0.84 (0.03) | 0.84 (0.03) | 0.83 (0.04) | 0.65 (0.06) |
| L-cau+acc | 0.89 (0.03) | 0.89 (0.03) | 0.90 (0.04) | 0.87 (0.02) |
| R-cau+acc | 0.84 (0.03) | 0.84 (0.03) | 0.83 (0.04) | 0.65 (0.06) |
| Combined mean | 0.89 | 0.87 | 0.88 | 0.77 |

Comparison of Dice score coefficient (DSC) mean values between manual and automated segmentation techniques. L – left, R – right, SD – standard deviation, GP- Globus Pallidus, Cau+acc – score calculated with the combined area of the caudate and the nucleus accumbens, Combined mean – mean score of all structures.

### Analysis of edits that were performed during manual segmentation

The edits were documented on 40 randomly chosen subjects of the total 80 to describe the workflow and also to highlight important areas for quality control. The hippocampus and amygdala consistently required the most edits. The hippocampus had two typical errors that required major manual corrections in most subjects: The lateral anterior superior border was overestimated in 35 and 36 subjects in the left and right hippocampus, respectively, and the inferior posterior area was too large in 30 and 32 subjects in the left and right hippocampus, respectively. The amygdala needed major edits on all subjects. The lateral superior border was overestimated in all subjects and the anterior side was underestimated in 33 and 35 subjects, for the left and right amygdala, respectively. The lateral inferior edge was too large in 21 on the left side and 18 on the right side. The thalami were overall slightly too big and needed minor edits throughout the structure, most notably on the medial posterior inferior edge, which was overestimated in 21 subjects for the left and in 19 for the right thalamus. The caudate received most edits on the lateral posterior inferior area, where the *FSL None* pipeline overestimated the border in 30 subjects for the left and in 26 for the right caudate. Notably the superior medial area of the right caudate was too large in 17 subjects, while on the left it was only overestimated in 3 subjects. All common edits are listed in **Table 4**. The putamen, GP and nucleus accumbens were more accurately segmented by FSL-FIRST than by FreeSurfer and only received minor and sporadic edits.

**Table 4.**
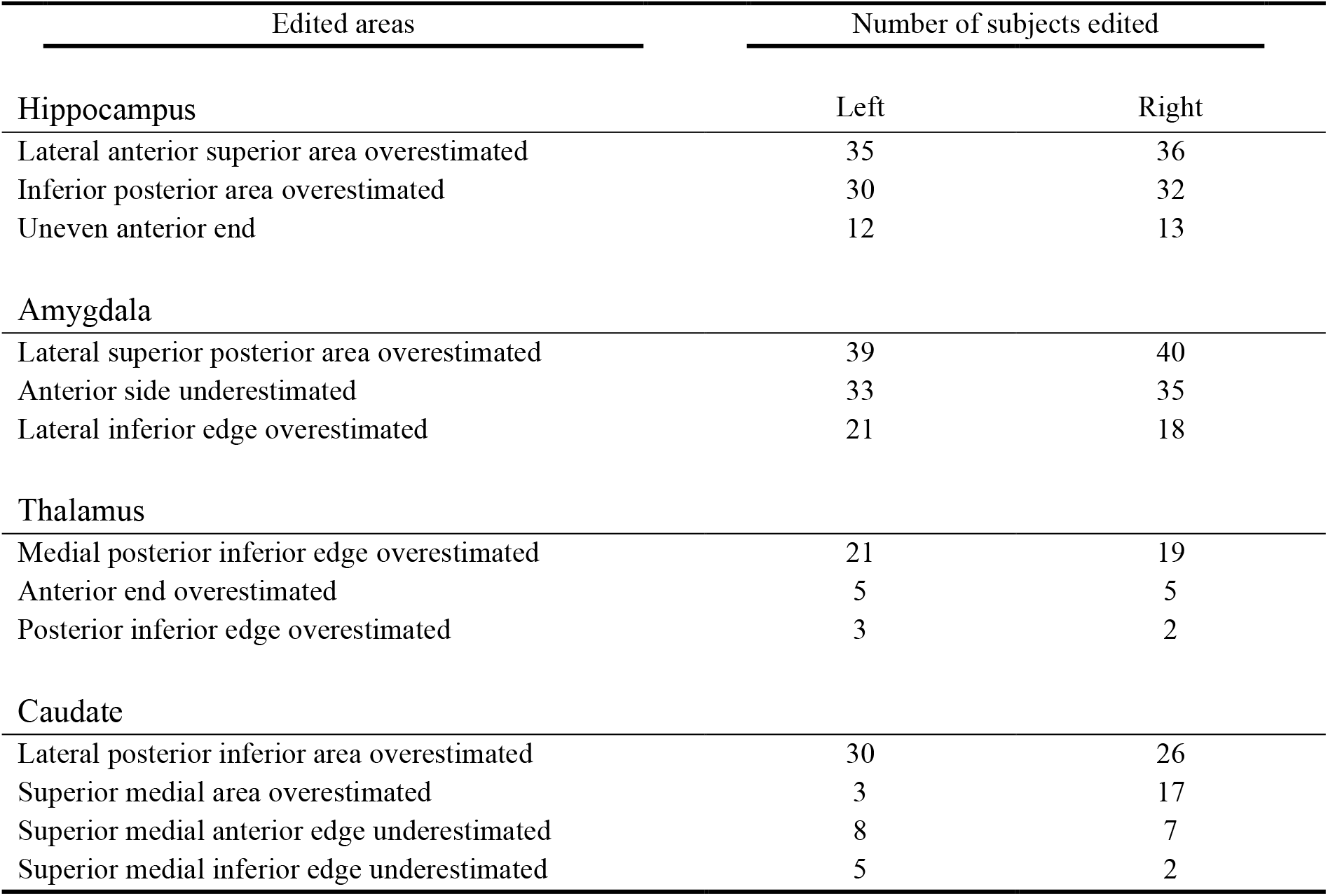
Most common major edits to structures and areas using the *FSL-None* segmentations out of 40 randomly chosen images.

| Edited areas | Number of subjects edited |  |
| --- | --- | --- |
|  | Left | Right |
| **Hippocampus** |  |  |
| Lateral anterior superior area overestimated | 35 | 36 |
| Inferior posterior area overestimated | 30 | 32 |
| Uneven anterior end | 12 | 13 |
| **Amygdala** |  |  |
| Lateral superior posterior area overestimated | 39 | 40 |
| Anterior side underestimated | 33 | 35 |
| Lateral inferior edge overestimated | 21 | 18 |
| **Thalamus** |  |  |
| Medial posterior inferior edge overestimated | 21 | 19 |
| Anterior end overestimated | 5 | 5 |
| Posterior inferior edge overestimated | 3 | 2 |
| **Caudate** |  |  |
| Lateral posterior inferior area overestimated | 30 | 26 |
| Superior medial area overestimated | 3 | 17 |
| Superior medial anterior edge underestimated | 8 | 7 |
| Superior medial inferior edge underestimated | 5 | 2 |

## Discussion

In this study we compared two automated segmentation tools, FSL-FIRST and FreeSurfer, against manual segmentation, on subcortical areas in a paediatric population. We included in the comparisons, FSL-FIRST’s three different pipelines, *FSL Default, FSL Fast* and *FSL None*, each of which uses different boundary correction settings to determine the exact anatomical borders of structures. Our goal was to compare the accuracy of these automated segmentation methods to manual segmentation, which is currently considered the gold standard (Hashempour et al., 2019; Morey et al., 2009), and has been validated as such in previous articles in paediatric as well as adult populations (Schoemaker et al., 2016; Makowski et al., 2018). In our results, *FSL Default* and *FSL Fast* pipelines performed overall more accurately than *FSL None* or FreeSurfer. We observed that automated methods tend to overestimate volumes in most structures, as was expected based on previous studies (Grimm et al., 2015; Hashempour et al., 2019; Nugent et al., 2013; Pipitone et al., 2014). The overestimation was overall most prominent with FreeSurfer and *FSL None*, although there were some notable exceptions in specific structures, such as the caudate, where FreeSurfer slightly underestimated volumes. Excluding the *FSL None* pipeline, FSL-FIRST produced generally better agreement across the structures than FreeSurfer.

### Hippocampus and amygdala

Both hippocampus and amygdala were overestimated by all automated segmentation methods in our study. Most accurate were *FSL Default* and *FSL Fast* pipelines with a moderate overestimation. *FSL None* and FreeSurfer overestimated both structures greatly. With all methods, the overestimation was more prominent in the amygdala than the hippocampus, which has also been documented in previous articles in adults as well as paediatric populations (Akudjedu et al. 2018; Doring et al., 2011; Pipitone et al., 2014; Schoemaker et al., 2016).

*FSL Default* and *FSL Fast* had overall better correlations with manual segmentation than *FSL None* or FreeSurfer. For the hippocampus, all of FSL-FIRST’s pipelines exceeded the threshold coefficient of r > 0.70, which has previously been suggested as the minimum for defining reliability between measures (Terwee et al., 2007). The Pearson correlation coefficients for the amygdala were lower, ranging from r = 0.61 to r = 0.67 with FSL-FIRST’s pipelines. FreeSurfers correlations were significantly weaker than FSL-FIRST’s for both hippocampus and amygdala, with amygdala having the lowest values. *FSL Default* and *FSL Fast* produced identical intraclass correlation (A, k) values, while *FSL None* and FreeSurfer showed very low to no correlation, indicating a large estimation bias. Automated segmentation of the hippocampus tends to have better consistency and reproducibility than the amygdala, which has been shown in multiple previous studies (Morey et al., 2009; Nugent et al., 2013; Pardoe et al., 2009; Schoemaker et al., 2016) that reported Pearson correlation coefficients ranging from r = 0.47 to r = 0.67 for the hippocampus and r = 0.24 to r = 0.35 for the amygdala using FSL-FIRST and r = 0.67 to r = 0.82 and r = 0.45 to r = 0.61 for the hippocampus and amygdala, respectively, using FreeSurfer. Similar results were shown regarding the DSC with every automated method producing higher mean values for the hippocampus (DSC > 0.76) than the amygdala (DSC > 0.60) in our results. The studies conducted by Morey et al. and Pardoe et al. also included DSC analysis showing results of the hippocampus producing higher spatial overlap than the amygdala with both FSL-FIRST and FreeSurfer, which is in line with our findings.

We found that FreeSurfer performed poorer than FSL-FIRST overall. This was an unexpected finding, as FreeSurfer has previously been reported to be overall more accurate and consistent than FSL-FIRST for both the hippocampus and amygdala for paediatric and adult populations (Morey et al., 2009; Schoemaker et al., 2016). Inter-rater variability may have contributed to these differences, as it is one of the key challenges with manual segmentation. The differences can be more pronounced in structures such as the amygdala, where the border around the structure may be difficult to distinguish visually. In these instances, the rater must rely on general anatomical knowledge instead of the intensities of the MR image to determine the exact shape of the structure. This is even more significant in paediatric MR images, since they have different contrast and comparatively lower resolution than adult images (Gousias et al., 2012). Example segmentations of the hippocampus and amygdala are presented in **Figure 2**.

**Figure 2.**
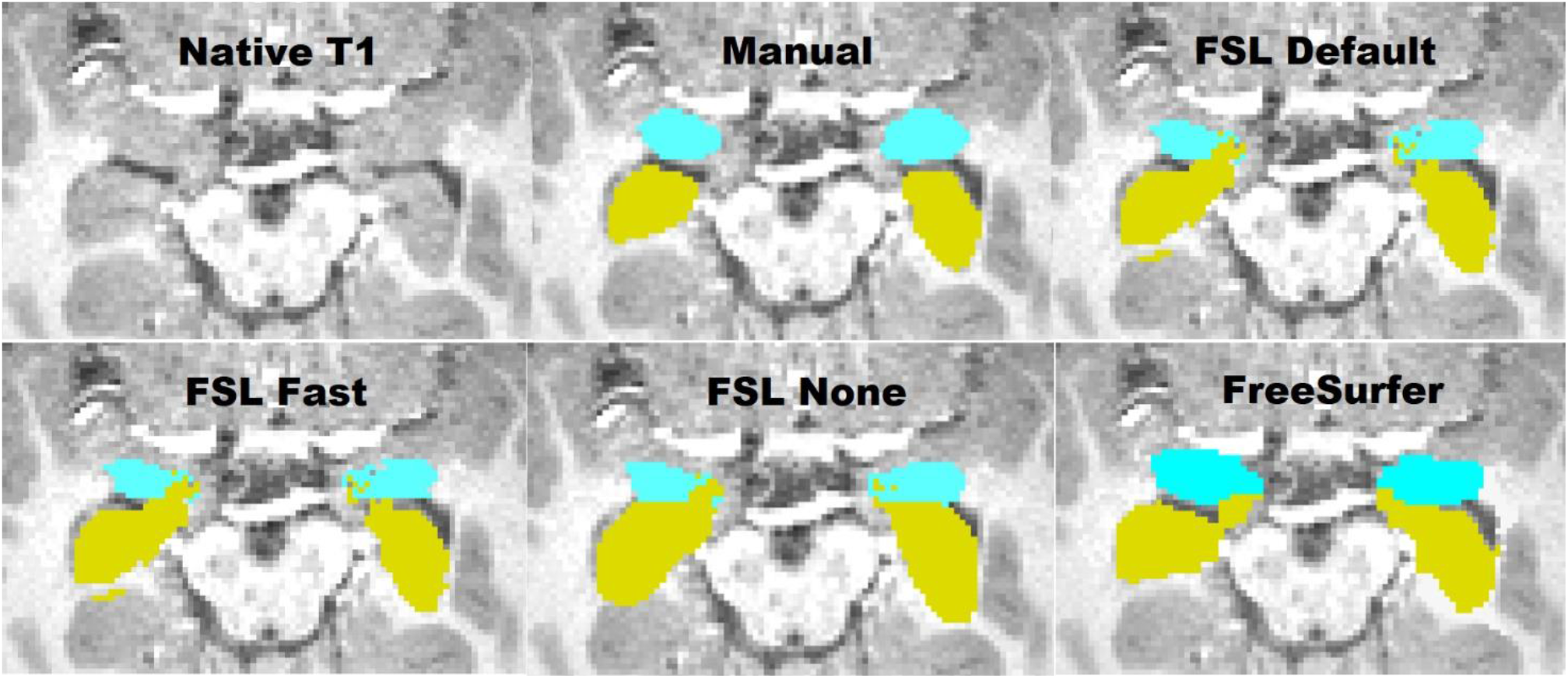
Transversal view of the segmentations of the hippocampus and amygdala. Yellow – hippocampus, turquoise – amygdala.

### Thalamus

The thalamus was most accurately segmented by FreeSurfer with only a slight overestimation. *FSL Default* and *FSL None* pipelines produced a larger overestimation while Fast underestimated the volume. Previous studies have shown results of FreeSurfer producing larger or similar volumes compared to FSL-FIRST (Hannoun, 2019; Makowski et al., 2018; Næss-schmidt et al., 2016). The discrepancy in results might be partly caused by inter-rater variability between the researchers in different studies. Despite having the most accurate mean volume, FreeSurfers Pearson correlation coefficient was significantly worse, r = 0.60, than any of FSL-FIRST’s pipelines, ranging from r = 0.86 to r = 0.89, indicating a larger volumetric variation in individual segmentations. Intraclass correlation (A, k) was on similar levels with coefficients ranging from ICC = 0.66 to ICC = 0.72, with all methods, suggesting a low to moderate reproducibility rate with manual segmentation. One previous study (Makowski et al., 2018) also showed weaker Pearson correlations for both FreeSurfer and FSL-FIRST than our results, ranging from r = 0.37 to r = 0.44, but included a significantly smaller sample size of 30 adults and that may explain some of the differences. The DSC values were great for all methods in our study, DSC > 0.91 for FSL-FIRST and DSC > 0.88 for FreeSurfer. A previous study done by Hannoun et al., including subjects aged between 1 and 18 years, showed similar results with DSC = 0.86 for FSL-FIRST and DSC = 0.84 for FreeSurfer (Hannoun et al., 2019). Segmentations of the thalamus are presented in **Figure 3** and **Figure 4**.

**Figure 3.**
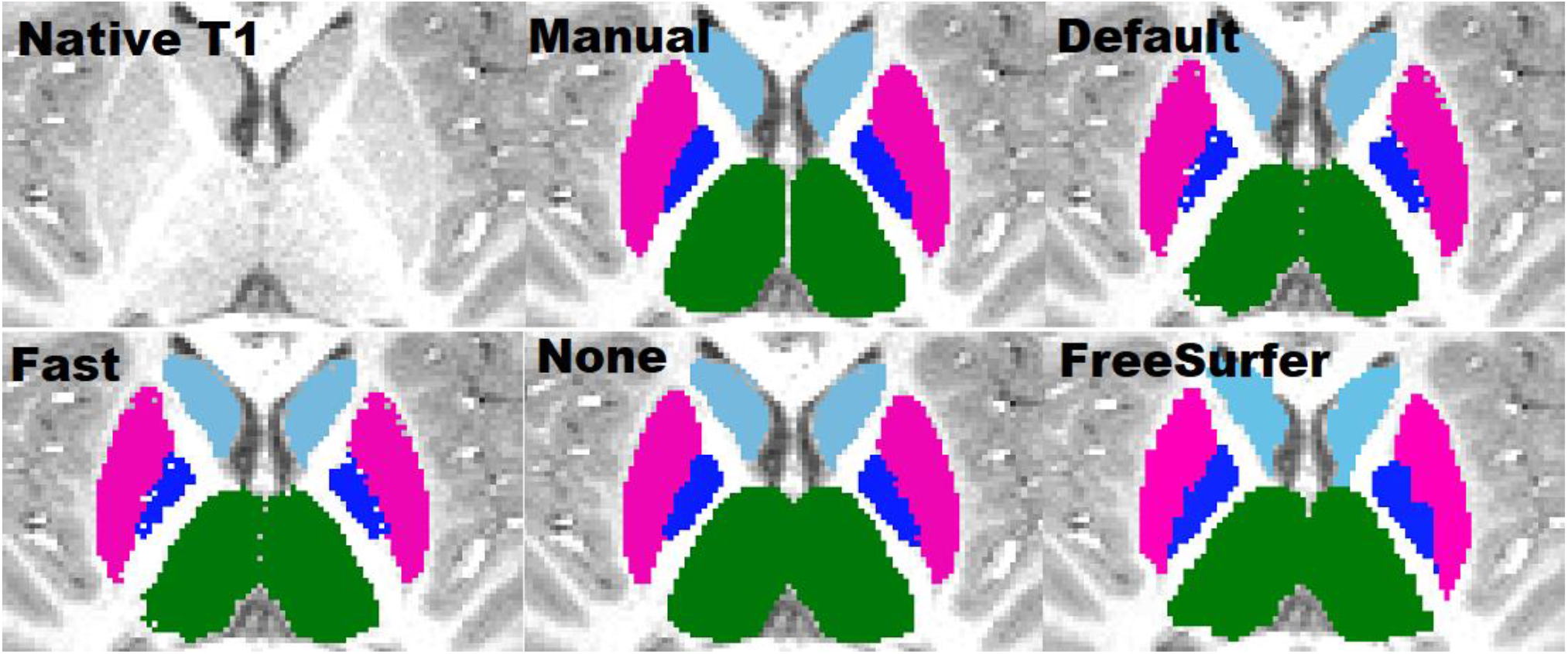
Transversal view of segmentations of the putamen, globus pallidus (GP), thalamus and caudate. Putamen – pink, GP – blue, thalamus – green, caudate – light blue.

**Figure 4.**
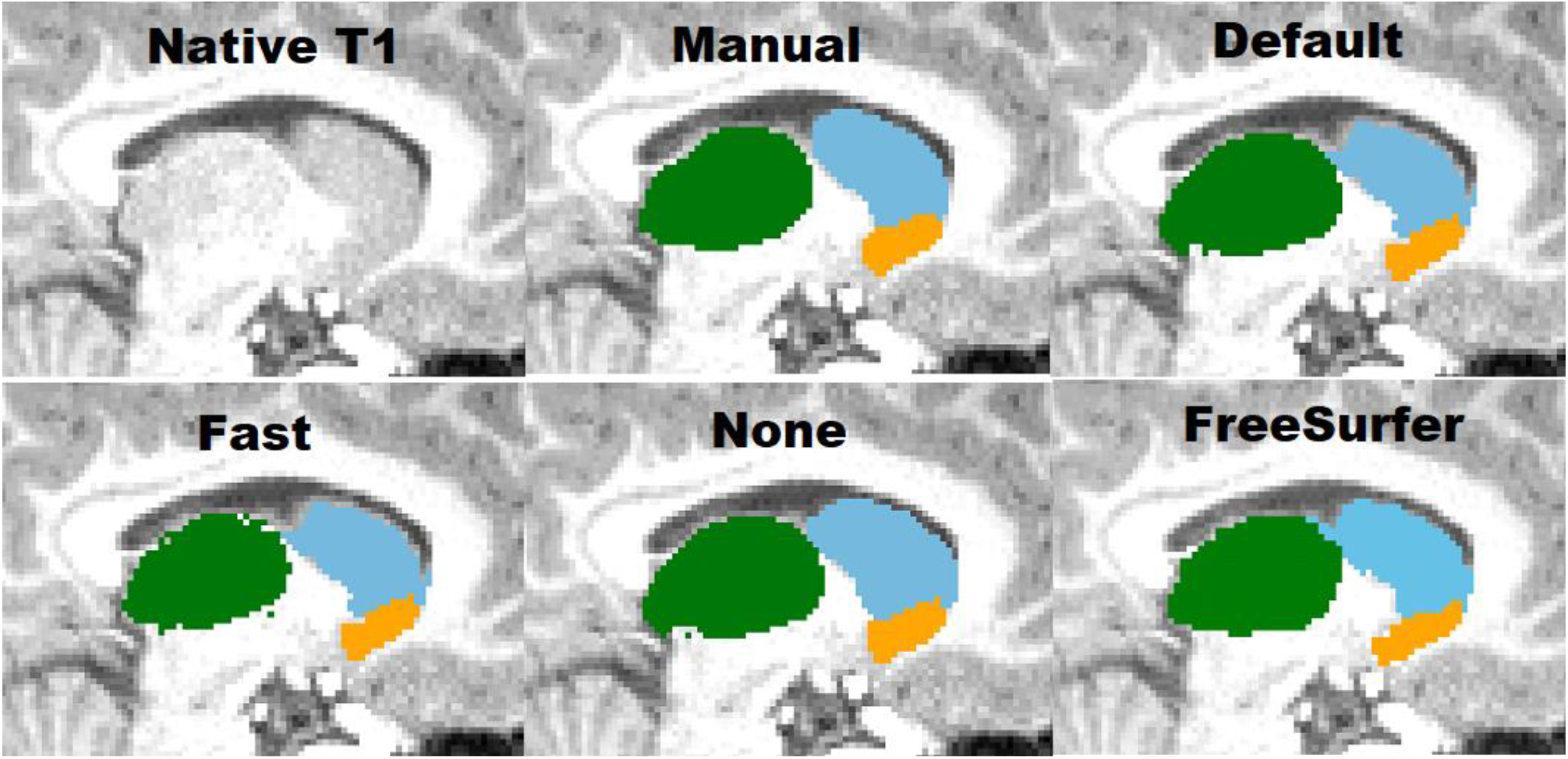
Sagittal view of the thalamus, caudate and nucleus accumbens. Thalamus – green, caudate – light blue, nucleus accumbens – orange.

### Putamen and globus pallidus

The putamen was segmented more accurately than the GP by all methods in this study. *FSL Default* and *FSL None* as well as FreeSurfer overestimated the putamen slightly, while *Fast* produced an underestimation of a similar volume. Similar results were observed with the GP, but with a greater magnitude. A previous study yielded similar results with FreeSurfer producing a higher overestimations than FIRST and GP having a greater relative volume difference than the putamen (Velasco-Annis et al., 2018). FSL-FIRST had excellent correlations for both putamen and GP, ranging from r = 0.86 to r = 0.98 across all pipelines. FreeSurfer also had a strong correlation for the putamen but performed significantly weaker for the GP with coefficients of r = 0.49 and r = 0.52 for the left and right GP. Intraclass correlation coefficients (A, k) were high across the board, with all methods yielding a coefficient of ICC > 0.80 for the putamen. For the GP, intraclass correlations were significantly lower for *FSL Fast* and FreeSurfer, while *FSL Default* and *FSL None* had great values of ICC > 0.80 for both structures, indicating a small estimation bias and good reproducibility with manual segmentation. A 2017 published study showed FreeSurfer having slightly better segmentation reproducibility for both the putamen and GP (Velasco-Annis et al., 2018). Another study published in 2018 showed the opposite and indicated that for FSL-FIRST has better consistency for the GP segmentation (Makowski et al., 2018). Direct comparison of these results is not ideal since both studies were done on an adult population and included a sample size of 30 or less. The DSC results in our study were great across the board with FSL-FIRST producing excellent results of DSC > 0.90 for both the putamen and GP with all techniques. FreeSurfer’s results were lower, but still satisfactory, DSC > 0.79. A previous study showed similar results with FSL-FIRST (DSC > 0.90) producing slightly higher DSC values than FreeSurfer (DSC > 0.80) for the putamen (Perlaki et al., 2017). However, the age of the subjects was not specified so the results may not be adequately comparable with our findings. To our knowledge this is the first automated segmentation method validation study done on a paediatric population including the putamen and GP. Segmentations of the putamen and GP are presented in **Figure 3**.

### Caudate and nucleus accumbens

The caudate was overall segmented accurately whereas the nucleus accumbens was overestimated by all methods in our study. The caudate was segmented accurately by all methods excluding *FSL None*, which overestimated both the caudate and the nucleus accumbens significantly. FreeSurfer and FSL-FIRST’s other pipelines produced an accurate volume for the caudate with only a minor underestimation. The nucleus accumbens was overestimated by all methods, with *FSL None* and FreeSurfer yielding the highest volumes. Notable is also the more prominent overestimation of the right nucleus accumbens, compared to the left, which was present in all four automated methods. Previous research indicates a moderate overestimation of both the caudate and nucleus accumbens with both FSL-FIRST and FreeSurfer (Perlaki et al., 2017; Velasco-Annis et al., 2018) with similar volumetric values compared to our results.

Pearson correlations coefficients were strong across all methods for the caudate, ranging from r = 0.69 to r = 0.84, showing a strong relationship between manual segmentation and the automated methods. The nucleus accumbens has similar coefficient values regarding FSL-FIRST, but FreeSurfer produced significantly weaker correlations. The intraclass correlation coefficients (A, k) showed that *FSL Default* and *FSL Fast* had superior reproducibility compared to *FSL None* and FreeSurfer for the nucleus accumbens. The results are similar for the caudate with the exception of FreeSurfer performing just as good as *FSL Default* and *FSL Fast*, with ICC values ranging from ICC = 0.85 to ICC = 0.90, while *FSL None’s* coefficients were significantly lower at ICC = 0.37 and ICC = 0.53 for the left and right caudate, respectively. The consistency and reproducibility of the caudate and nucleus accumbens have been documented in previous studies with slightly different results compared to our study (Perlaki et al., 2017; Velasco-Annis et al., 2018). The article by Velasco-Annis et al. suggested great reproducibility rates for the caudate with both FreeSurfer and FSL-FIRST, with ICC values ranging from ICC = 0.86 to ICC = 0.93, producing similar values for each method. The other study conducted by Perlaki et al. showed a slightly better reproducibility with FreeSurfer regarding the caudate and nucleus accumbens. The study by Perlaki et al. also showed results similar to ours regarding the DSC values with FSL-FIRST producing better slightly better values than FreeSurfer for the caudate (Perlaki et al., 2017).

Overall, these variations in results may be explained with the difficult determination of the border between the caudate and nucleus accumbens. The intensities of the MR image are visually indistinguishable for these two structures, which may lead to inaccuracy in volumetric quantification. To assess this problem, we combined the volumes of both structures to eliminate possible errors caused by the similarity of intensities. Considering the relatively small volume of the nucleus accumbens, the results for combined volume were similar to the results derived from the caudate volumes. Segmentations of the caudate and nucleus accumbens are presented in **Figure 4**.

### Limitations

Our study presents a few limitations. Firstly, the sample size is limited due to the time-consuming manual segmentation process but likely sufficient for building study-specific templates, which is a potential goal for applied studies (Lee et al., 2019). Secondly, all manual segmentations were performed by a single rater which might lead to some systematic biases in delineation of anatomical borders in MR images. However, the expert review provides some safeguard for this. On a related note, the manual segmentation was done by editing models produced by *FSL None* which might potentially cause the manual segmentations to have a bias towards FSL-FIRST.

## Conclusions

In this feasibility study, we determined the accuracy of two automated segmentation tools for T1-weighted MR images, FSL-FIRST with three different boundary correction settings and FreeSurfer against manual segmentation in a paediatric, 5-year-old population (N = 80). Overall, the automated tools show promising accuracies, but the performance of all automated tools changed vastly based on the structure. Small structures such as the amygdala and nucleus accumbens were inaccurately segmented by all automated methods. On the other hand, the segmentation of the putamen and the caudate were performed accurately with most of the automated methods and yielded relatively good consistency and reproducibility with manual segmentation. The use of these automated segmentation tools in neuroimaging studies still presents challenges, and careful visual inspection of the automated segmentations is still strongly advised, since there are many factors, such as the quality of the used MR-images that might impact the accuracy of the segmentations. Future research should investigate the benefits of using custom subcortical atlases to improve the accuracy and reliability of automated segmentation methods especially for the amygdala and hippocampus (Lee et al., 2019).

## Acknowledgements

The authors declare no conflict of interest.

EPP was supported by the Päivikki and Sakari Sohlberg Foundation. ESi was supported by Juho Vainio Foundation, Finnish Brain Foundation, Turunmaan Duodecim-seura. VK was supported by Finnish Cultural Foundation, Lastenlinnan säätiö. NH was supported by the Orion Research Foundation, the University of Turku Graduate School and the Hospital District of Southwest Finland State Research Grants. LK was supported by Brain and Behaviour Foundation, NARSAD, YI Grant #1956, State Grants for Clinical Research (ERVA), the Academy of Finland (Profi 5, #325292) SN was supported by the State Grants for Clinical Research. JJT was supported by the Hospital District of Southwest Finland, Turku University Foundation, State Grants for Clinical Research and Emil Aaltonen Foundation, Alfred Kordellin Foundation (data collection and data analysis) as well as Sigrid Jusélius Foundation (interpretation of the data and writing the manuscript).

